## Supplementary Information and Figures for "Multimodal analysis of RNA sequencing data powers discovery of complex trait genetics"

#### Implementation

The backbone of our Pantry implementation (https://github.com/daniel-munro/Pantry) is a pipeline built with Snakemake. Snakemake is a workflow management system with an emphasis on bioinformatics pipelines. It provides flexibility both in terms of data and project parameters, which can be specified in configuration files, and in terms of computing environments, such as high-performance computing clusters.

The code for the data processing pipeline consists of existing programs, e.g., STAR and samtools, additional scripts to process their input and output data, and the Snakemake code that runs the programs and scripts. All of this pipeline code is contained within a project template directory within Pantry that can be copied, customized, and executed. This allows the pipeline to be customized in terms of parameters or the addition of entirely new phenotypes, while preserving the customized code alongside the data it produces. A second project template directory with similar structure holds the Pheast downstream genetic analysis module.

The default Pantry pipeline was designed for computational and storage efficiency. For example, several modalities involve read quantification with respect to modality-specific annotations. Instead of using a quantification method that requires separate read alignments for each set of annotations, Pantry uses kallisto, a pseudoalignment method, for quantification, reducing runtime and greatly reducing intermediate file storage. Similarly, for the BAM alignment files used for other modalities to count intron junctions, exon reads, and intron reads, unused read information is removed from the BAM files, reducing their size by an order of magnitude without affecting the results.

Pantry includes a module called Pheast (PHEnotype Application STreamlined) for running various downstream analyses with all Pantry-generated phenotypes. As with the phenotype-generating code, Pheast is packaged as its own project directory template with a configuration file, Snakemake code, and scripts. It can be copied and edited to run analyses with generated Pantry phenotypes and corresponding genotypes. This two-stage approach allows the complexities of phenotype generation, including reference files, properties of the input sequence data, and the variety of phenotyping software, to be of concern only in the first stage. Then, in the second stage, multiple phenotype sets, formatted uniformly as BED files and accompanying metadata, can be used for each of multiple downstream applications. By default, this stage includes generation of covariates from genotypes and the phenotypes in each set, followed by cis-QTL mapping and fitting xTWAS (FUSION) models.

### Supplementary Figures

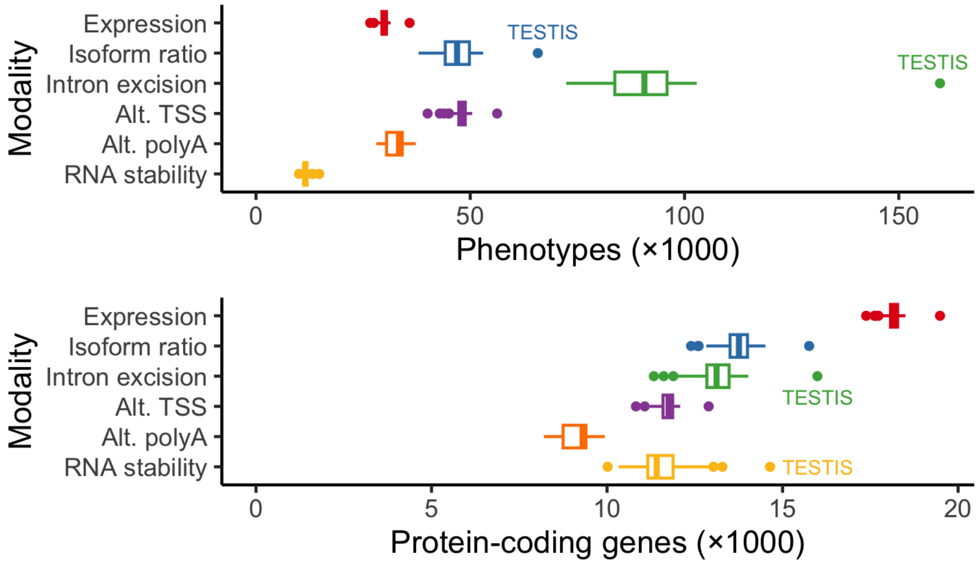


**Figure S1.** Above: boxplots showing the number of phenotypes extracted per GTEx tissue per modality, for all protein-coding genes and lncRNAs. Below: the number of protein-coding genes represented by the phenotypes above.

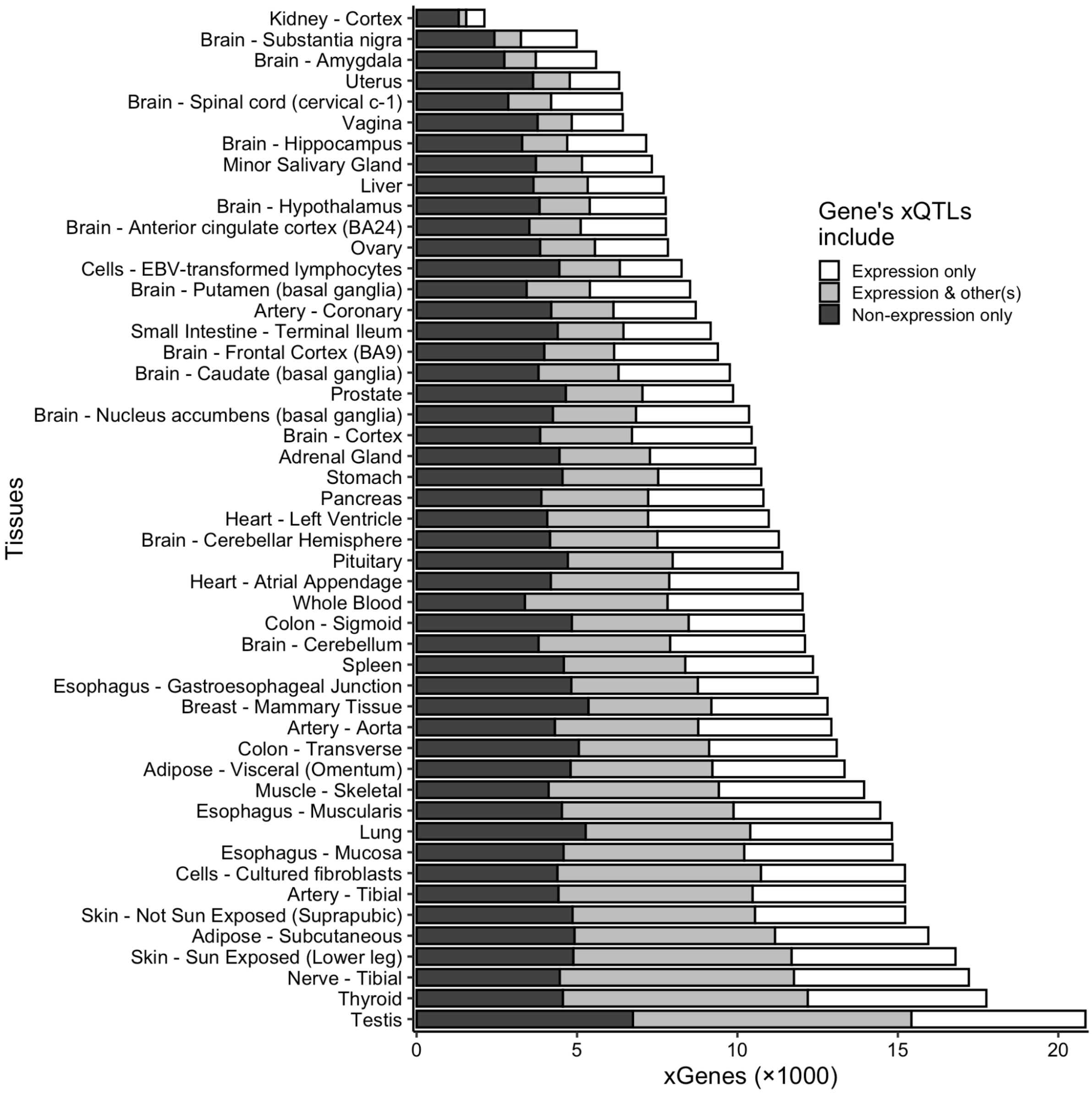


**Figure S2.** For each GTEx tissue, the number of genes with at least one xQTL are shown, colored by whether each gene's xQTL(s) included eQTLs, one or more other modality of xQTL, or both. Results are shown for xQTLs mapped separately per modality to show the increase in gene count compared to eQTL mapping alone.

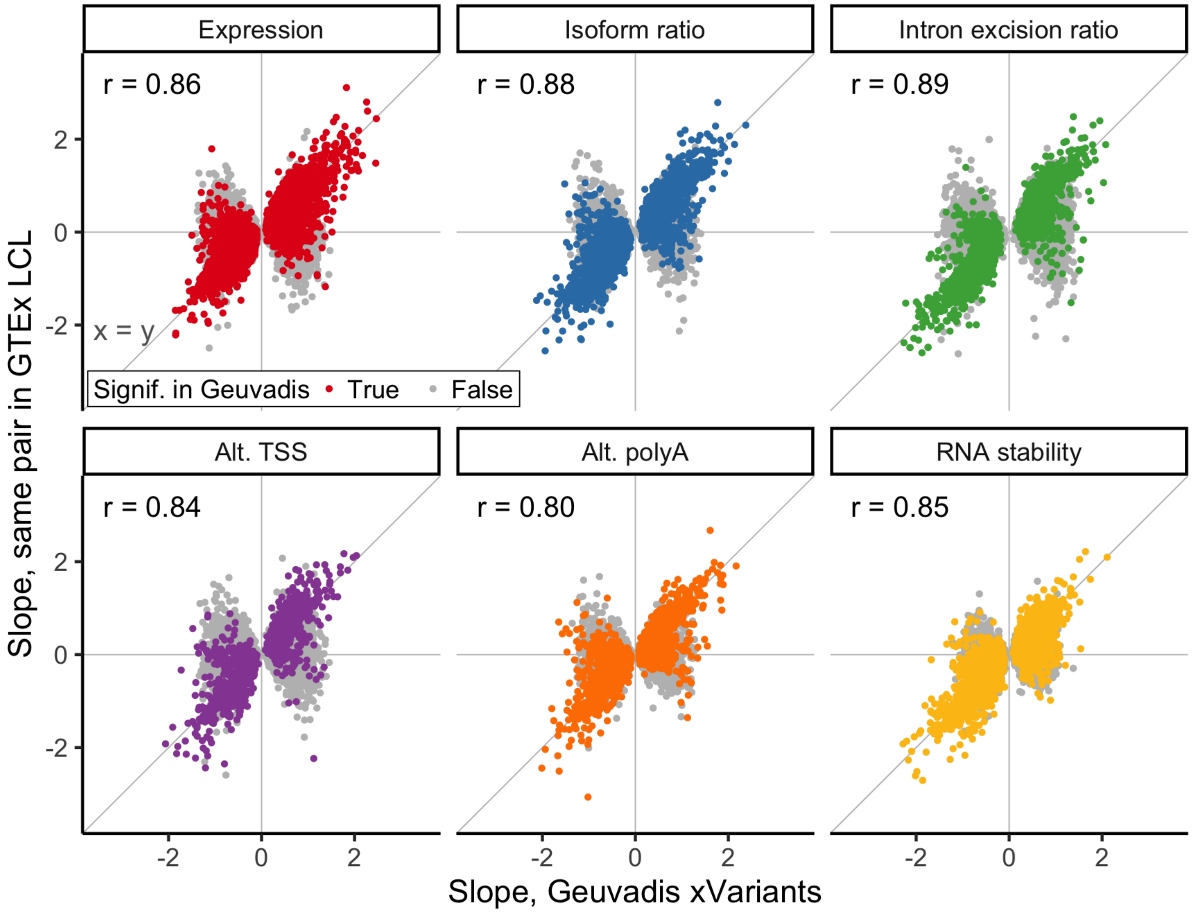


**Figure S3.** Concordance of xQTL associations between independent datasets. For each modality, we identified the top xQTL association per gene in Geuvadis and extracted the slope of the same variant-phenotype pairs, if tested, from GTEx LCL tissue results. Associations that were significant in Geuvadis are colored according to modality, and nonsignificant associations are gray. Pearson's correlation coefficient was calculated for the significant associations per modality.

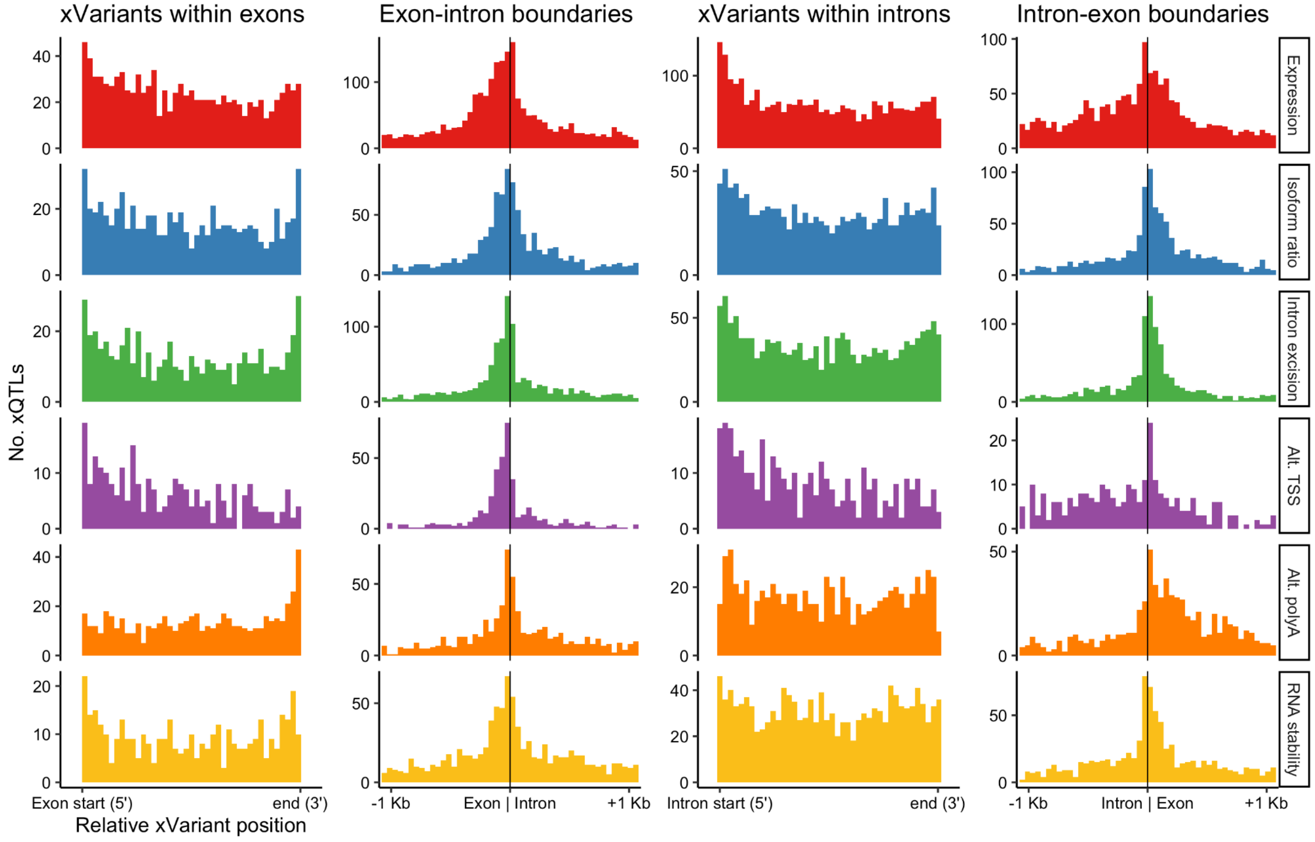


**Figure S4.** Location of xQTL variants from combined-modality mapping in Geuvadis, relative to exons, introns, and boundaries. For the first and third columns, the genomic coordinates of xVariants were linearly transformed such that the exon/intron starts and ends were aligned on the x-axis. The second and fourth columns show the distribution of QTL positions within 1 Kb of all exon-intron and intron-exon boundaries, oriented 5’-3’, without normalizing by gene length.

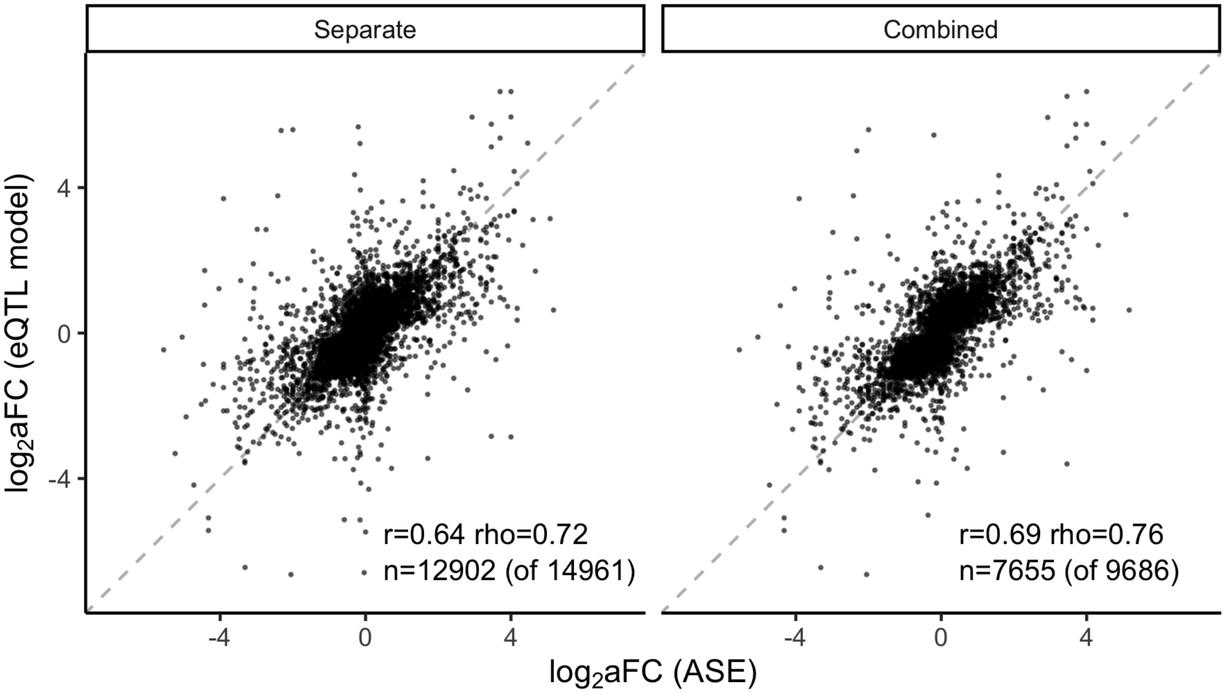


**Figure S5.** Scatter plots of allelic fold change of eQTLs in subcutaneous adipose GTEx tissue (ADPSBQ), when mapping conditionally independent xQTLs for modalities separately (left) and together (right). aFC was measured for each eQTL gene-variant pair using aFC-n (i.e., the eQTL model) and using allele-specific expression (ASE) in heterozygous individuals. Pearson's r and Spearman's rho are shown for both eQTL sets, along with the number of gene-variant pairs included, which are the subsets of eQTLs for which aFC could be measured from ASE.

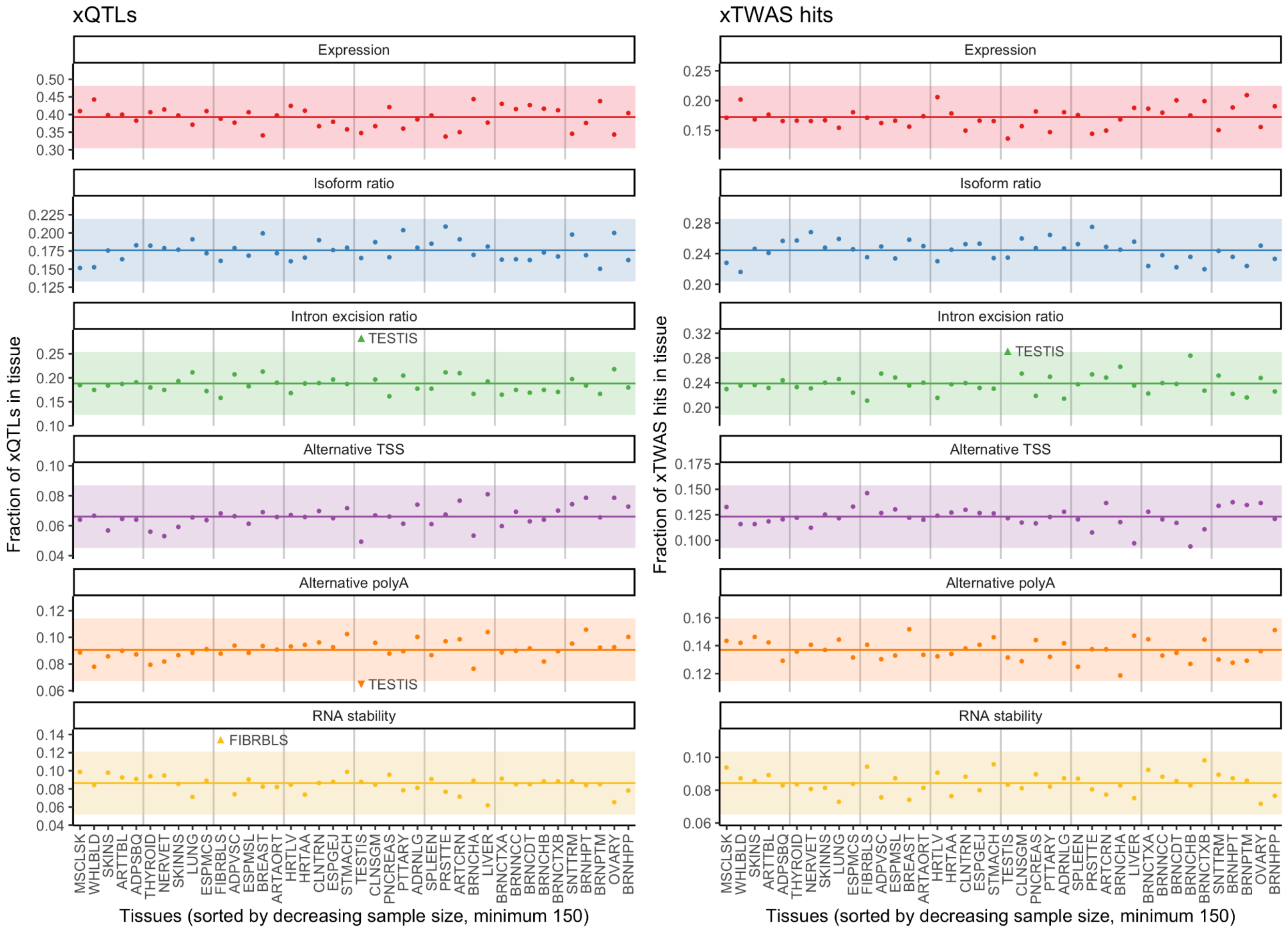


**Figure S6**. Proportions of RNA modalities in cis-QTLs and TWAS hits. Left: Fraction of cis-QTLs per modality in each GTEx tissue with sample size >150. Horizontal lines and bands indicate mean and three standard deviations per modality. Upward and downward-pointing triangles indicate values above and below the ±3 standard deviation interval, respectively. Right: The same plot but for fraction of TWAS hits, across all 114 tested traits.
